# The Mediterranean mussel, *Mytilus galloprovincialis*, a novel model for developmental studies of mollusks

**DOI:** 10.1101/2023.07.27.550798

**Authors:** Angelica Miglioli, Marion Tredez, Manon Boosten, Camille Sant, João E. Carvalho, Philippe Dru, Laura Canesi, Michael Schubert, Rémi Dumollard

**Author notes:** Co-First authors (Angelica Miglioli, Marion Tredez). Co-Last authors (Michael Schubert, Rémi Dumollard).

## Abstract

A model organism in developmental biology is defined by its experimental amenability as well as by resources created for the model system by the scientific community. For the most powerful models, the combination of both has already yielded a thorough understanding of development. However, the number of developmental model systems is still very limited, and their phylogenetic distribution is heavily biased. Members of one of the largest animal phyla, the mollusks, for example, have long been neglected as developmental model organisms. To remedy this shortcoming, we produced a detailed developmental transcriptome for the Mediterranean mussel *Mytilus galloprovincialis*, a bivalve mollusk, and expanded the list of experimental protocols available for this species. Our high-quality transcriptome allowed us to identify transcriptomic signatures of developmental transitions and to perform a first comparison with the Pacific oyster *Crassostrea gigas* that can be used in future multi-species analyses. To allow co-labelling studies, we optimized protocols for immunohistochemistry and hybridization chain reaction and combined both techniques to create high-resolution co-expression maps of developmental genes. The resources and protocols we describe here thus represent an enormous boost for the establishment of the Mediterranean mussel as a laboratory model in developmental biology.

**Summary statement:** Resources and techniques are described for the Mediterranean mussel *Mytilus galloprovincialis*, which, together, establish a novel model system for studying mollusk development and animal evolution.

## 1. Introduction

The development of aquatic invertebrates has inspired the curiosity of biologists for well over a century (De Robertis and Tejeda-Muñoz, 2022; Hunt and Maslakova, 2017; Nielsen, 2005, 2009; Wang et al., 2020). Morphological similarities between larvae have, for example, been used as taxonomic characters and comparisons between larval types have yielded highly influential hypotheses on animal evolution (Carrier et al., 2018). One such larval type, the trochophore, is widely distributed amongst the lophotrochozoans and may have already been present in the last common ancestor of this large clade of bilaterian animals (De Robertis and Tejeda-Muñoz, 2022). However, despite decades of studies (Carrier et al., 2018; Carrillo-Baltodano et al., 2021; Nielsen, 2005; Page, 2009; Rawlinson, 2010; Wada et al., 2020; Xu et al., 2016; Yang et al., 2020), our understanding of the evolutionary origin of the trochophore larva as well as of the biological processes underpinning its ontogeny and transition to the adult body remains extremely fragmentary (Carrier et al., 2018; De Robertis and Tejeda-Muñoz, 2022; Wada et al., 2020; Wang et al., 2020; Wu et al., 2019; Xu et al., 2016; Yang et al., 2020). Although multiple genome and transcriptome sequencing projects have provided abundant data from diverse organisms, most of these datasets have yet to be exploited to provide insights into the overall conservation of trochophore development between different taxa (Liu et al., 2014; Sayers et al., 2021; Wada et al., 2020). A major limitation for comparisons seems to be that lophotrochozoan development is characterized by the deployment of species-specific gene sets. This fact makes the identification of comparable larval phases, even between closely related species of the same class, quite difficult (Wada et al., 2020; Wu et al., 2019; Xu et al., 2016; Yang et al., 2020).

Within the lophotrochozoans, the mollusks are the largest and anatomically most diverse phylum (Yang et al., 2020). Yet, as of today, the scientific community still lacks an amenable laboratory model for studying the mechanisms of mollusk development. Reasons include the limited availability of genomic and transcriptomic resources and the complexity of establishing even basic techniques, such as immunocytochemistry and *in situ* hybridization (Yang et al., 2020). It has been proposed that the overall fragility of mollusk larvae as well as the presence of a shell, at least in conchiferan (i.e., shell-bearing) mollusks (Kocot et al., 2016; Miglioli et al., 2019; Miglioli et al., 2021a; Sayers et al., 2021; Wada et al., 2020; Yang et al., 2020), have hindered efforts to devise more elaborate resources and techniques for representatives of this phylum (Carrier et al., 2018; Sayers et al., 2021; Wada et al., 2020). Altogether, these issues have slowed research into the molecular aspects of trochophore development in mollusks, hence limiting our capacity for comparisons of trochophore development between mollusks and other taxa (Wada et al., 2020; Yang et al., 2020).

The Mediterranean mussel, *Mytilus galloprovincialis*, is an established model in environmental studies, and a species of great ecological and economic interest with a worldwide distribution (Brzozowska et al., 2012; Daguin and Borsa, 2000; FAO, 2022). Large-scale larval cultures of *M. galloprovincialis* are easily set up in the laboratory, following standardized culture protocols by the International Standardization Organization (ISO) and the American Society for Testing Materials (ASTM) (ASTM E724-21, 2021; ISO 17244:2015, 2020). *M. galloprovincialis* is also the first metazoan with a published pangenome (Gerdol et al., 2020), a feature of significant importance for researchers interested in genome evolution. The presence of detailed culture protocols and of a well-annotated genome make *M. galloprovincialis* a promising model organism for developmental studies. In addition, basic protocols for pharmacological treatments, quantification of shell biogenesis, quantitative polymerase chain reaction (qPCR) as well as immunohistochemistry and *in situ* hybridization have recently been described for developing *M. galloprovincialis* embryos and larvae (Miglioli et al., 2019, 2021a, 2021b).

In this work, we provide a compendium of updated protocols and novel resources to significantly facilitate access to and work with *M. galloprovincialis* embryos and larvae. We thus compiled a detailed developmental transcriptome and validated it with a standardized qPCR protocol. We further established a novel protocol for multi-color, fluorescent whole-mount *in situ* hybridization, using the hybridization chain reaction (HCR) approach (Choi et al., 2014), and successfully coupled this *in situ* hybridization approach with immunohistochemistry assays. The developmental transcriptome of *M. galloprovincialis* allowed us to define major developmental periods and transitions, results that were further confirmed in other bivalve mollusks and that provide a basis for future comparative developmental studies of lophotrochozoans. The novel *in situ* hybridization and immunohistochemistry protocols yielded detailed information on the expression of genes marking developmental transitions, for example, in the early shell field, apical sensory cells, peripheral mantle cells, and hindgut. Taken together, this work represents a valuable resource for future work with *M. galloprovincialis* embryos and larvae, setting the stage for establishing this animal as a mollusk model system. As such, this work will have a significant impact on developmental biology and associated research fields.

## 2. Results

### 2.1. Morphological staging of *Mytilus galloprovincialis* early development and description of transcriptome libraries

The developmental transcriptome of *M. galloprovincialis* embryos and larvae consists of 15 developmental stages (*i.e.*, a total of 30 libraries from two independent biological replicates). Samples were obtained every 4 hours under standard culture conditions at 16°C (ASTM E724-21, 2021; ISO 17244:2015, 2020), from unfertilized egg to 52 hours post fertilization (hpf) with an additional sampling timepoint at 72 hpf (Fig. 1A). As previously described in other Mytilids, the *M. galloprovincialis* zygote undergoes holoblastic, unequal cleavage (Kurita et al., 2009). The micromeres form at the animal pole of the embryo at 4 hpf, and subsequent cleavage follows a spiral pattern (Kurita et al., 2009). By 12 hpf, the macromeres at the vegetal pole start to invaginate, indicating the start of gastrulation. At 16 hpf, the stomodeum (or presumptive mouth) has moved anteriorly, and a rosette of dorsal ectodermal cells forms an invagination, indicating the onset of shell field ontogeny (Kniprath, 1980, 1981; Miglioli et al., 2019). At 20 hpf, the columnar cells of the shell field thicken, creating the impression of a deeper invagination. At 24 hpf, the larvae have the stereotypical morphology of a trochophore, with a ventral stomodeum and a dorsal shell field that is now completing the evagination process and secreting the organic matrix of the larval shell (Kniprath, 1980; Miglioli et al., 2019). By 28 hpf, the hinge region of the shell field is completely flat. At 32 hpf, a cavity appears in the hinge region of the larva, and, at 36 hpf, it expands and seems to connect to the former stomodeum (that has now moved anteriorly) and to the anal cavity on the posterior side. At 40 hpf, the D-margin of the shell becomes clearly visible and only the velum (also called the prototroch) of the larva is protruding. By 44 hpf, the velum is enclosed in the D-shaped shell. Thereafter, no major morphological changes are observable, excepting the progressive outgrowth of the shell with respect to the mantle tissue along the D-border (Fig. 1A). In summary, our transcriptome is representative of all developmental stages of early bivalve development.

**Figure 1:**
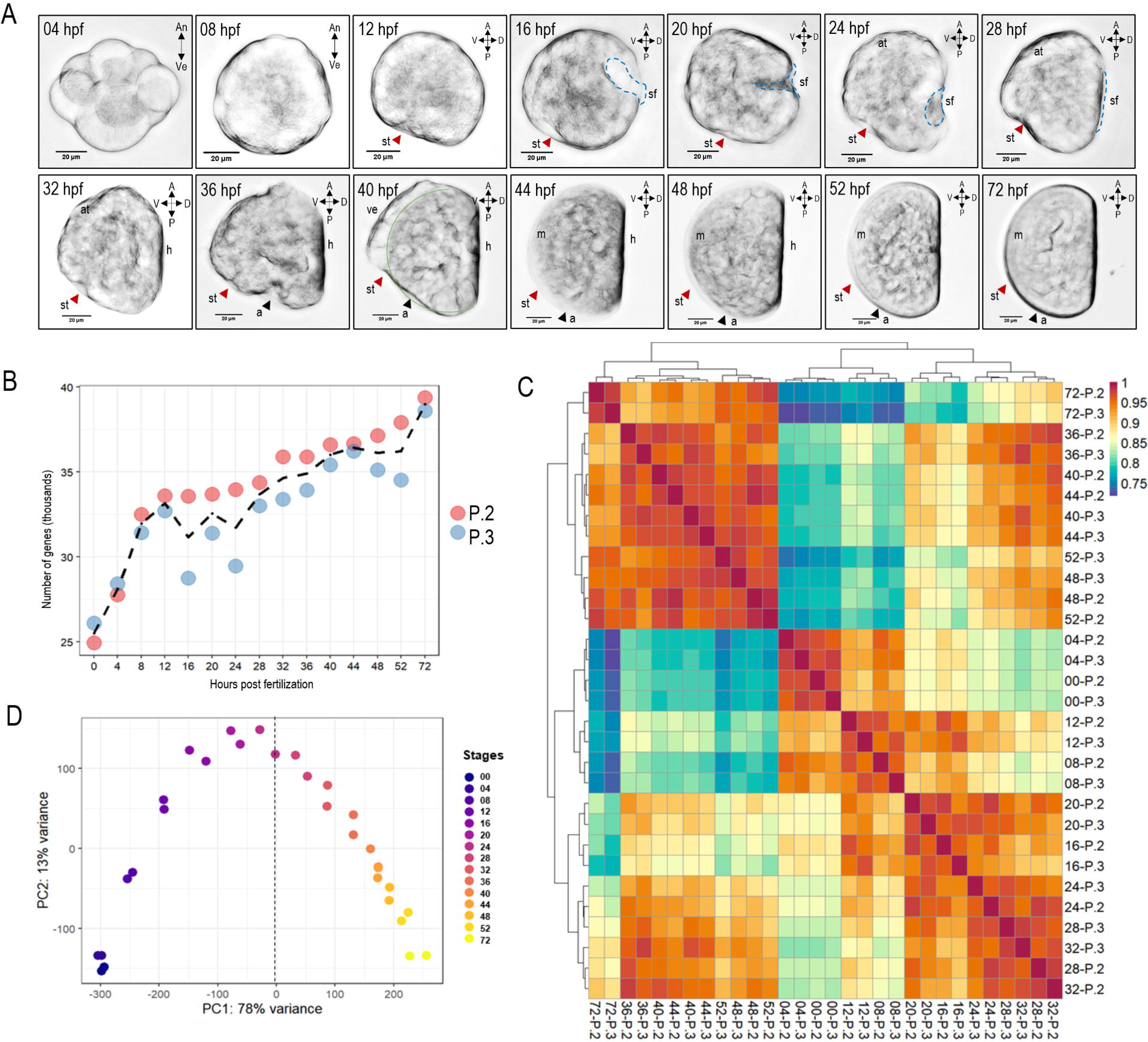
Morphological and transcriptomic staging of *Mytilus galloprovincialis* early development. A) Development of *M. galloprovincialis* under standard culture conditions. Orientation is as shown by arrows in the top right of each picture, hours post fertilization (hpf) (at 16°C) is indicated on the top left. Red and black arrowheads highlight stomodeum/presumptive mouth and anus, respectively, blue dotted line indicates the shell field outline (16 to 24 hpf) and the hinge region (28 hpf). At 40 hpf, the green line indicates the D-margin of the calcified shell. Abbreviations: An: animal; Ve: vegetal; A: anterior; P: posterior; D: dorsal; V: ventral; a: anus; at: apical tuft; h: hinge; m: mantle; sf: shell field; st: stomodeum/presumptive mouth; ve: velum. Scale bar: 20 μm. B) Dot plot of the number of genes expressed in each library before DESeq2 normalization. The black dotted line represents the mean trend throughout development. C) Heatmap of the Euclidean distance matrix of the libraries clustered with default hclust parameters. D) Principal component analysis (PCA) of DESeq2 normalized libraries, dotted line indicates the axis of principal component 2.

### 2.2. Transcriptome assembly and gene expression dynamics

More than 31 million paired-end clean reads per library were mapped on the published *M. galloprovincialis* genome (Gerdol et al., 2020), with an average input length of 200 bp (Supplementary Table 1). More than 85% of the reads were mapped, with 71% of them assigned to a unique locus (Supplementary Table 1). The output gene count matrix included 52,422 expressed genes (sum of gene raw counts in all libraries > 0) of a total of 60,302 entries in all libraries (Supplementary Table 2). An analysis of the number of genes expressed during development is presented in Fig. 1B. The minimum number of expressed genes (EGs) was found in unfertilized eggs (at 0 hpf), with around 25,000, while the highest was found at the late D-veliger stage (at 72 hpf), with around 40,000, indicating that at least 15,000 genes (that is about 25% of the genome) are transcriptionally modulated during early development of *M. galloprovincialis* (Fig. 1B). The number of EGs increased dramatically during the first 12 hours of development, with the two replicates yielding similar results. Thereafter, a slow but steady increase of EGs was observed until the last sampling point. A heatmap of the Euclidean distances between stages and a principal component analysis (PCA) of the normalized libraries validated sample reproducibility (Fig. 1C and 1D). The heatmap further revealed that the libraries clustered by developmental time and not by biological replicate (Fig. 1C). Library clustering yielded two main clusters, one before and one after 32 hpf, indicating a potential mid-development transcriptomic shift within the trochophore stage (Fig. 1C). The PCA showed a continuous transcriptomic progression from 0 to 72 hpf along the axis of the first principal component, indicating that the maximum variation is found between the 0 to 24 hpf and the 32 to 72 hpf libraries (Fig. 1D), which is consistent with a potential transcriptomic shift during early trochophore development. A majority of the variance was explained by the first principal component (78%), which is mainly correlated with developmental timing, lending support to the notion that morphological changes during development are driven by transcriptomic changes.

### 2.3. Differential expression analysis and developmental clustering

Differential expression analysis was performed by pairwise comparison of samples of consecutive stages. The analysis identified 11,996 differentially expressed genes (DEGs) during *M. galloprovincialis* development (Supplementary File 2 and Supplementary Table 3). Hierarchical clustering of the libraries based on DEG expression values identified statistically distinct developmental phases (adjusted-unbiased p-values, au > 0.95) (Fig. 2A) and divided the 30 libraries in two main clusters: one regrouping the libraries from 0 to 16 hpf (embryonic phase) and one regrouping the libraries from 20 to 72 hpf (larval phase). In the embryonic phase of development, the 0 and 4 hpf libraries were clearly distinct from the ones covering 8 to 16 hpf, corresponding to, respectively, zygote and gastrulation (Fig. 2A). In the larval phase of development, three sub-clusters were observable. The first one was formed by libraries corresponding to three trochophore stages, from 20 to 28 hpf, with three older trochophore-like larvae (advanced trochophore), from 32 to 40 hpf, establishing a second cluster. The third cluster grouped the libraries of the four stages with D-veliger morphology, from 44 to 72 hpf. The number of DEGs varied as development proceeded, with an overall decreasing trend (Fig. 2B). About 55% of DEGs were found to be exclusively up-regulated, 25% exclusively down-regulated, and the remaining 20% alternated between up- and down-regulation. Of note, 40% of DEGs were differentially expressed more than once during development (Fig. 2B, insert). More than 1,500 DEGs were already detected at 4 hpf, indicating an early initiation of zygotic gene expression. The highest number of DEGs was reached at 8 hpf, and it was followed by a progressive decrease as development proceeded. Following the differential expression analysis, DEGs were grouped as a function of their most recurrent expression profiles (Fig. 2C and 2D, Supplementary Tables 4 and 5). The analysis identified 56 clusters with more than 15 genes and of the top 6 clusters, based on the number of DEGs, we found 5 to consist of genes with low maternal (0 hpf) expression levels. This indicates that the DEGs we identified are predominantly derived from zygotic expression (Fig. 2C). An expression level-based association of gene clusters with developmental periods (Supplementary Table 4) showed that 31% of the clusters reached their highest expression levels at the zygote, 32% at the gastrula, 14% at the trochophore, 7% at the advanced trochophore, and 25% at the D-veliger stage (Fig. 2D). These results indicate that zygote and D-veliger stages are characterized by genes with a higher diversity of expression profiles than the trochophore and advanced trochophore stages. Moreover, the analysis indicated that expression levels of individual clusters are generally skewed towards earlier developmental periods and that cluster expression levels might be correlated with key morphological events during development of both embryo and larva.

**Figure 2:**
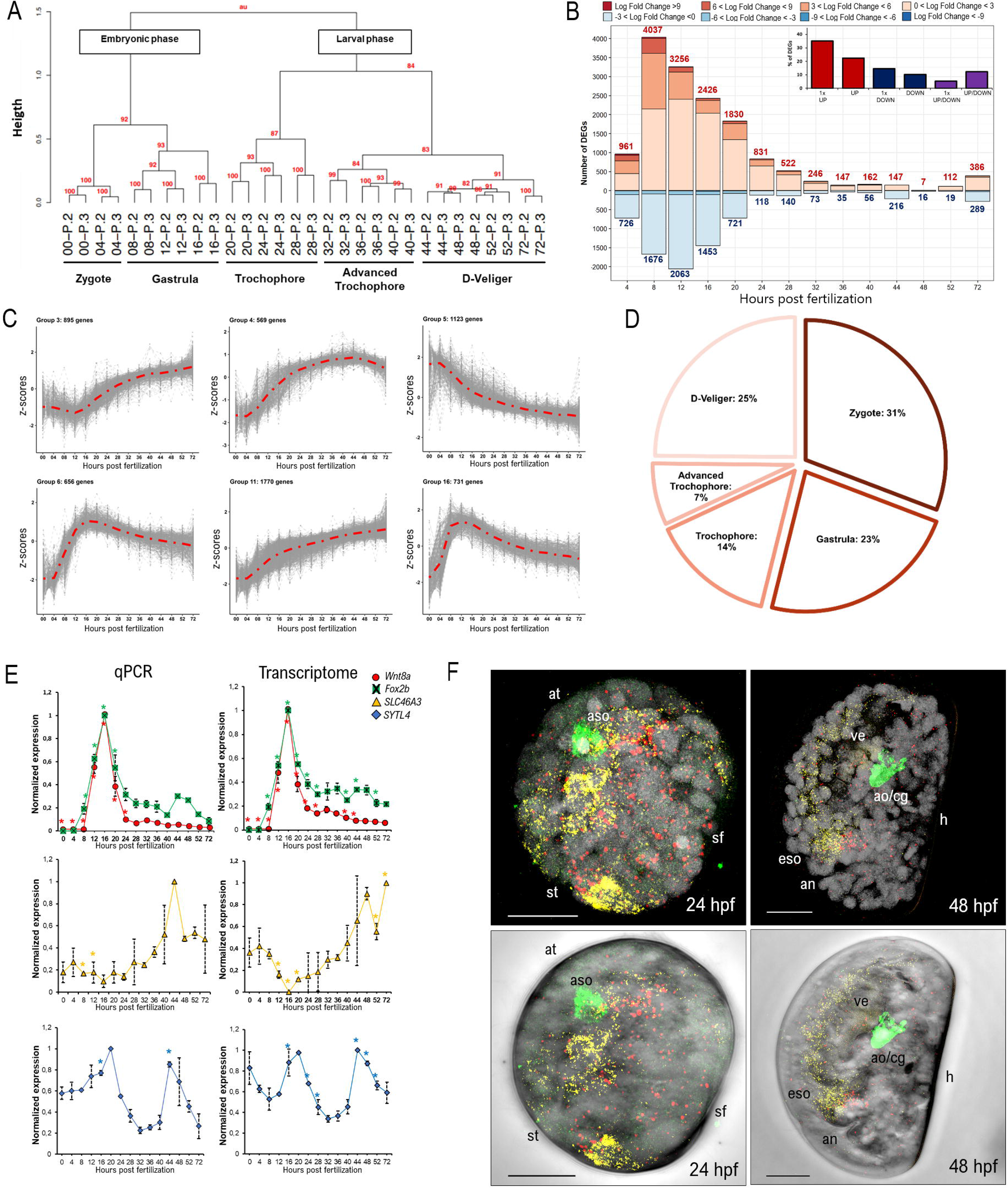
Differential expression analysis highlights stage-specific transcriptomic signatures. A) Hierarchical clustering of differentially expressed gene expression values in the transcriptome libraries. Bootstrap values are shown in red on each branch. Names for each cluster are assigned according to morphology and developmental timepoint. B) Histogram showing the number of up- (red) and down- (blue) regulated genes at each assayed stage. Color code is according to log fold change values. Insert: percentage (%) of differentially expressed genes that are up-, down- or up-/down-regulated in the dataset, either once (1x) or multiple times. C) Top six gene expression profiles by number of genes. D) Pie graph depicting the percentage (%) of profiles reaching their highest expression levels at different developmental stages. E) Normalized expression of *Wnt8a*, *Foxb2*, *SLC46A3*, and *SYTL4* from transcriptomic (left) and qPCR (right) data. F) Maximum projection superimpositions of *Foxb2* (yellow) and *Wnt8a* (red) fluorescent *in situ* hybridization with serotonin (5-HT) immunohistochemistry (green) and Hoechst (grey) or in bright field in trochophore (24 hpf) and D-veliger (48 hpf) larvae. Abbreviations: an: anus; ao/cg: apical organ/cerebral ganglion; at: apical tuft; aso: apical sensory organ; eso: esophagus; h: hinge; sf: shell field; st: stomodeum/presumptive mouth; ve: velum. Scale bar: 20 μm.

### 2.4. qPCR and *in situ* hybridization of selected genes

The differential expression analyses were corroborated by qPCR experiments carried out with material from both biological replicates. Four target genes were selected based on their transcriptomic expression profiles. The *Foxb2* (MGAL_10B093191) and *Wnt8a* (MGAL_10B085403) genes were selected as members of one of the top 6 clusters (Fig. 2C, Group 6), characterized, respectively, by high and low mean expression levels. *SLC46A3* (MGAL_10B020966) and *SYTL4* (MGAL_10B044489) were selected because they showed a highly dynamic expression profile (Supplementary Tables 3 and 4). For each one of the four genes, the qPCR experiments validated both the levels and the temporal patterns of their expression (Fig. 2E). Given their shared temporal expression patterns, we further assessed the spatiotemporal expression of *Foxb2* and *Wnt8a* in relation to each other and relative to the immunoreactivity of serotonin (5-HT), a marker of mussel nervous system cells. To do so, we established a fluorescent double *in situ* hybridization protocol based on the hybridization chain reaction (HCR) protocol for *M. galloprovincialis* and combined it with an immunohistochemistry assay for serotonin (Fig. 2E and Supplementary Figure 1). Co-labeling with serotonin allowed a standardized orientation of embryos and larvae, which significantly facilitated the interpretation of the *in situ* hybridization signals. We thus found that, while *Wnt8a* was detectable in a discrete group of cells that will give rise to one of the posterior neural ganglions of the larva, *Foxb2* was detectable in the stomodeal and trochal area. These observations were consistent with the expression patterns described for these two genes in other lophotrochozoan species (Lartillot et al., 2002; Marlow et al., 2014; Tomer et al., 2010). In sum, qPCR analysis and *in situ* hybridization confirmed the developmental trends identified by our transcriptome analyses.

### 2.5. Qualitative analysis of differentially expressed genes

We next performed a qualitative analysis of the DEGs based on functional domains and gene ontologies (GOs) (Fig. 3). DEGs were screened for transcription factor domains (Fig. 3A), domains known to be involved in shell formation (Fig. 3B) as well as for receptors, ion channels, and neurotransmitter transporters (Fig. 3C). In total, 711 transcription factors with 65 different domains were annotated, with the most recurrent (> 20 genes) being: zinc fingers, homeodomains, helix-loop-helix and fork head domains (Fig. 3A, Supplementary Table 6). With respect to shell formation, 779 genes were found to contain domains associated with shell matrix proteins and calcification (Fig. 3B) (Ramesh et al., 2019). The most recurrent domains included calcium binding proteins (EF-hand and calponin), von Willebrand factors (WKA), extracellular matrix proteins (fibronectin), carbohydrate binding proteins (lectin and concanavalin), chitin binding proteins (CB), ion channels, and transporters (Fig. 3B, Supplementary Table 5). Regarding receptors, 225 DEGs were annotated as G protein-coupled receptors of the rhodopsin and secretin families, 92 as ephrin-A receptors, 23 belonged to the nuclear receptor superfamily, and other DEGs included scavenger, low density (LD) lipoprotein, and TGFβII receptors (Fig. 3C). As many as 125 DEGs were predicted to be involved in neurotransmitter transport and reception (Fig. 3C, Supplementary Table 3). A gene ontology enrichment analysis was subsequently performed on the DEGs to identify stage-enriched biological processes (Fig. 3D). The GO enrichment analysis for biological processes identified 6 GO clusters: transmembrane transport, actin polymerization, cell adhesion, receptor-mediated signal transduction, biosynthesis, and transcription-coupled repair. While most biological processes did not exhibit a clear association with a specific developmental phase, transmembrane transport and receptor-mediated signal transduction were enriched in the trochophore and advanced trochophore stages (20 to 28 hpf and 32 to 40 hpf). Actin polymerization and transcription-coupled repair were weakly enriched in embryonic development (zygote and gastrula), and biosynthesis in the larval phase (from the early trochophore onward). Altogether, the results of the quantitative analysis confirmed the relevance of the identified DEGs with respect to the major regulatory (*i.e.*, transcription factors) and ontogenetic (*i.e.*, shell matrix proteins) programs controlling early development of bivalve mollusks.

**Figure 3:**
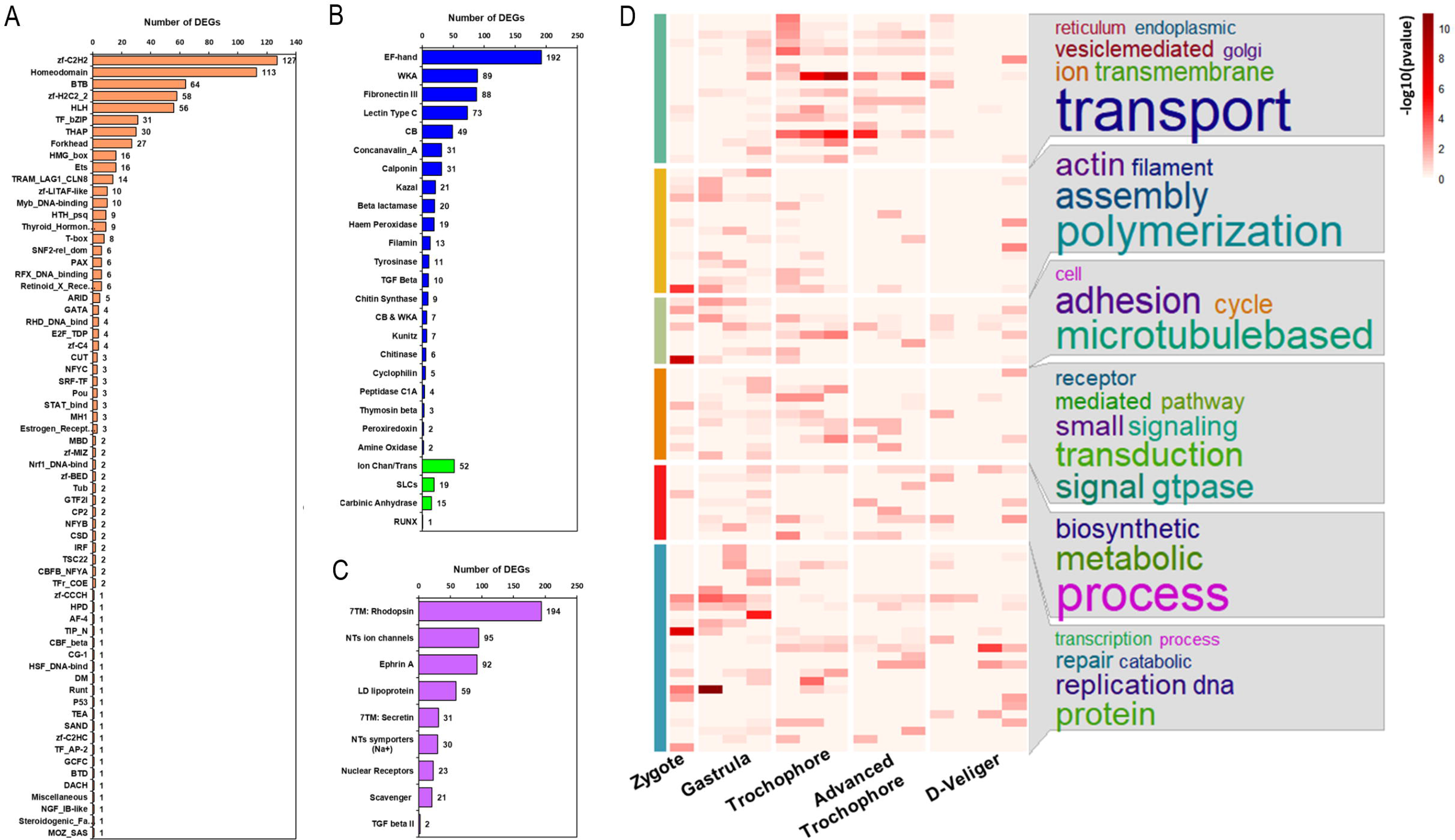
Qualitative analysis of functional domains and gene ontologies (GOs) of differentially expressed genes (DEGs). A) Transcription factors. B) Domains associated with shell biogenesis: shell matrix proteins in blue and calcification proteins in green. C) Receptors and neurotransmitter transporters. D) Composite image showing the heatmap of significant GO terms paired with GO cluster analysis at different stages of development. Colors indicate the -log_10_(p-value). Column breaks in the heatmap divide the different developmental stages identified by hierarchical clustering, and row breaks separate the GO clusters. Row color annotation (on the left) highlights different GO clusters, and the most significant terms for each cluster are shown in their respective box (on the right).

### 2.6. Gene clustering and transcriptomic signatures of early developmental transitions

Soft clustering of expressed gene was performed to reveal potential transcriptomic dynamics during development. The analysis identified nine soft clusters (Fig. 4A), which were named according to the developmental staging system described above (Fig. 1A). While the embryonic gene clusters nicely recovered the hierarchical clusters of the embryonic phase, the larval soft clusters predominantly highlighted stages between the developmental phases identified by hierarchical clustering, thereby revealing potential developmental transitions. The early trochophore gene cluster matched the hierarchical clusters transitioning from the gastrula to the trochophore (16 to 20 hpf, Transition 1), the advanced trochophore gene clusters matched the hierarchical clusters transitioning from trochophore to advanced trochophore (28 to 32 hpf, Transition 2), and the veliger gene cluster matched the hierarchical clusters transitioning from advanced trochophore to D-veliger (40 to 44 hpf, Transition 3) (Fig. 4A, Supplementary Table 7). To identify representative genes for developmental Transitions 1, 2, and 3, we isolated the genes of the three soft clusters that were differentially expressed at the corresponding transition stages (Supplementary Table 8). We found 98, 11, and 8 genes for, respectively, Transition 1, 2, and 3 (Fig. 4B, Supplementary Table 3). To assess whether similar transition signatures exist in other bivalves, we searched for homologous genes in the Pacific oyster *Crassostrea gigas* and analyzed their developmental expression using publicly available transcriptomic data (Fig. 4C, Supplementary Table 8) (Liu et al., 2021). The results of these comparisons indicated that, of the 98 Transition 1 genes in *M. galloprovincialis*, 22 were characterized by conspicuous expression at gastrula and trochophore 1 stages in *C. gigas*. However, most of these genes were also highly expressed at earlier or later developmental stages (Fig. 4C). Of the 11 *M. galloprovincialis* Transition 2 genes, we identified two characterized by high expression between the corresponding stages in *C. gigas*, *i.e.*, trochophore 3 and trochophore 4 (Fig. 4C). For Transition 3, marked by 8 genes in *M. galloprovincialis*, we found that three genes in *C. gigas* were characterized by a peak of expression at the passage from advanced trochophore (trochophore 5) to early veliger (early D-veliger 1) (Fig. 4C). Our unbiased cluster analysis thus revealed three developmental transitions during early *M. galloprovincialis* development that are impossible to identify morphologically and that are at least partially conserved in other bivalves.

**Figure 4:**
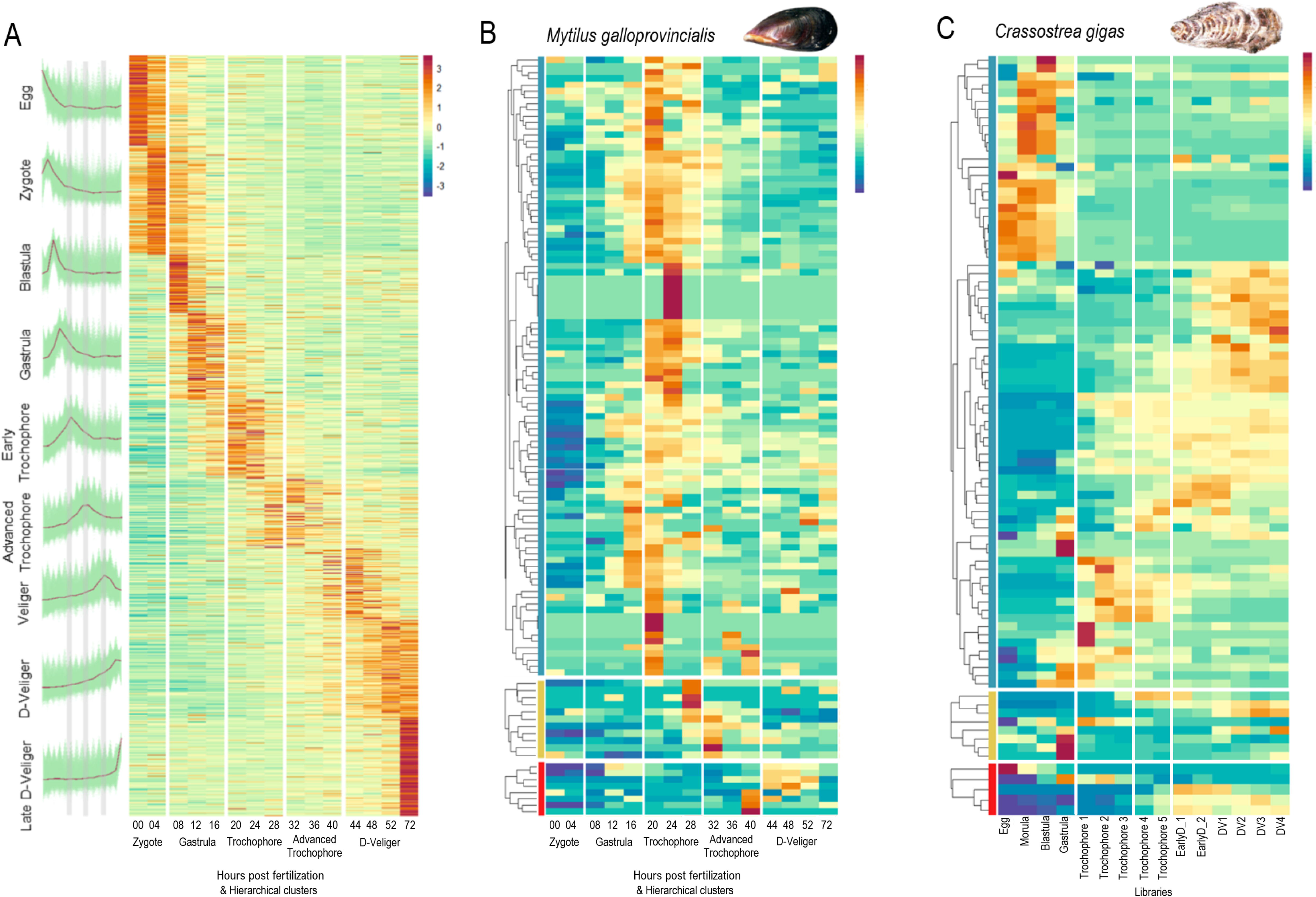
Soft clustering reveals transcriptomic signatures of early developmental transitions. A) Left panel: expression patterns of single soft clusters. The green lines depict the expression pattern in scaled transcripts per million (z-scores) of each gene belonging to a soft cluster, the red line indicates the mean trend of the expression data, vertical grey lines indicate developmental transitions. Right panel: heatmap of all soft clusters in scaled transcripts per million (z-scores) arranged by developmental time of appearance. White lines identify developmental transitions identified by hierarchical clustering. B) Expression of developmental transition genes in scaled transcripts per million (z-scores) in the mussel (*Mytilus galloprovincialis*) transcriptome. C) Expression of developmental transitions genes in scaled transcripts per million (z-scores) in the Pacific oyster (*Crassostrea gigas*) developmental transcriptome. Column breaks indicate the three transitions. Color annotations on the left, in B) and C), highlight the three developmental transitions: blue, gold, and red for Transition 1, 2, and 3, respectively.

### 2.7. Tissue localization of conserved transition genes by *in situ* hybridization

To shed light on the potential developmental functions of the conserved transition genes, we assessed the expression of a set of these genes by *in situ* hybridization using our HCR protocol. Genes were selected based on their differential expression in *M. galloprovincialis* (log fold change > 2) and their expression profiles in *C. gigas* (z-score value becoming positive) at the timepoints of the respective transition (Table 1, Supplementary Figure 2, Supplementary File 2). The genes were named according to the results of BLAST searches with their predicted proteins of the Uniprot and NCBI databases (Table 1, Supplementary Table 3). As Transition 1 contained a significantly higher number of conserved genes, we selected 8 genes for Transition 1 and two genes each for Transitions 2 and 3. Expression of these genes was analyzed at their corresponding transition stages. At 20 hpf, the transcription factor genes *Tbx2b* and *Erg* were detectable in cells surrounding the stomodeum and in the apical tuft, with *Tbx2b* also being expressed in the hindgut (Fig. 5A). The other 6 assayed genes of Transition 1 were all expressed in or in association with the shell field: *HPP*, *TYRO2*, and *Pxdn* were detectable in groups of cells surrounding the invaginating shell field at 16 hpf, with *TYRO2* maintaining this expression at 20 hpf, *Dmbt1* in column-like cells at 16 and 20 hpf, *NA03* in islets of cells that elongated from invaginated tissue at 20 hpf, and *SLC4A4* in layers of cells covering the invaginated tissue at 20 hpf (Fig. 5A). For the conserved Transition 2 genes, *SLC26A5* was detectable in two symmetric groups of ectodermal cells at 28 hpf (Fig. 5B). At that stage, *Grin3a* was weakly expressed throughout the post-trochal larva, with a small group of anterior cells beneath the apical tuft expressing the gene more conspicuously (Fig. 5B). The two selected genes of Transition 3 were both expressed symmetrically with respect to the hinge axis at 44hpf. Thus, while *EF-hand* was expressed in the velum, the esophagus, and in a dense group of cells close to the anus (Fig. 5C), *COLVI* was detectable in individual cells along the D-border of the mantle (Fig. 5C).

**Figure 5:**
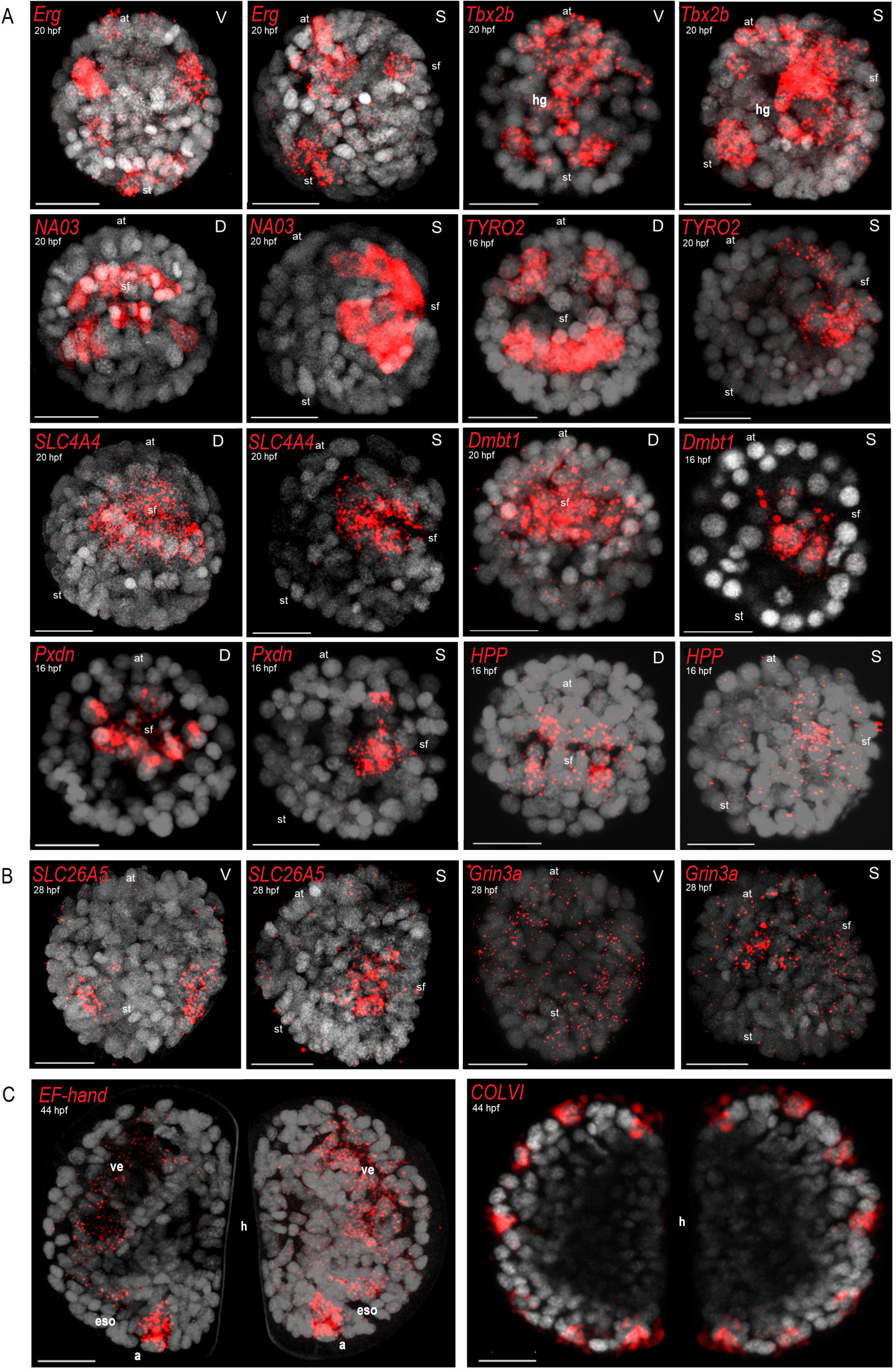
Developmental expression of homologous transition genes in *Mytilus galloprovincialis* larvae. A) Genes selected for Transition 1 (gastrula to trochophore). B) Genes selected for Transition 2 (trochophore to advanced trochophore). C) Genes selected for Transition 3 (advanced trochophore to D-veliger). Left and right valves are presented next to each other, relative to the hinge region. The *in situ* hybridization signal is shown in red, Hoechst-stained nuclei are shown in grey. Gene names are indicated in red in the top left corner. Developmental timing in hours post fertilization (hpf) (at 16°C) is indicated in white under the gene name. Abbreviations: D: dorsal view; S: sagittal view; V: ventral view; a: anus; at: apical tuft; eso: esophagus; h: hinge; sf: shell field; st: stomodeum/presumptive mouth. Scale bar: 20 μm.

**Table 1.**
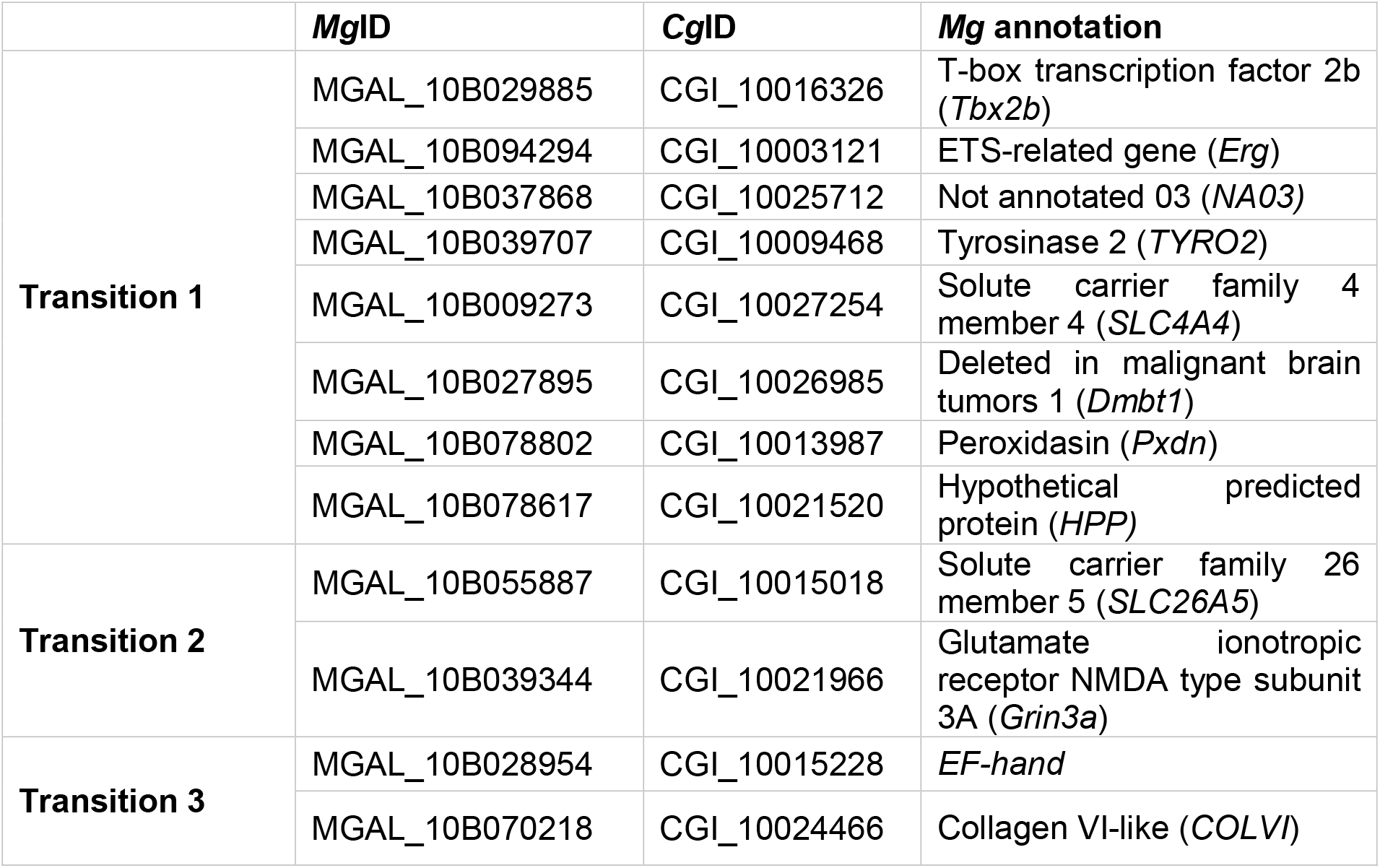
List of conserved transition genes in *Mytilus galloprovincialis* (*Mg*ID) and *Crassostrea gigas* (*Cg*ID) selected for *in situ* hybridization in *M. galloprovincialis*.

|  | **MgID** | **CgID** | **Mg annotation** |
| --- | --- | --- | --- |
| **Transition 1** | MGAL_10B029885 | CGI_10016326 | T-box transcription factor 2b (*Tbx2b*) |
|  | MGAL_10B094294 | CGI_10003121 | ETS-related gene (*Erg*) |
|  | MGAL_10B037868 | CGI_10025712 | Not annotated 03 (*NA03*) |
|  | MGAL_10B039707 | CGI_10009468 | Tyrosinase 2 (*TYRO2*) |
|  | MGAL_10B009273 | CGI_10027254 | Solute carrier family 4 member 4 (*SLC4A4*) |
|  | MGAL_10B027895 | CGI_10026985 | Deleted in malignant brain tumors 1 (*Dmbt1*) |
|  | MGAL_10B078802 | CGI_10013987 | Peroxidasin (*Pxdn*) |
|  | MGAL_10B078617 | CGI_10021520 | Hypothetical predicted protein (*HPP*) |
| **Transition 2** | MGAL_10B055887 | CGI_10015018 | Solute carrier family 26 member 5 (*SLC26A5*) |
|  | MGAL_10B039344 | CGI_10021966 | Glutamate ionotropic receptor NMDA type subunit 3A (*Grin3a*) |
| **Transition 3** | MGAL_10B028954 | CGI_10015228 | *EF-hand* |
|  | MGAL_10B070218 | CGI_10024466 | Collagen VI-like (*COLVI*) |

## 3. Discussion

### 3.1. Fine-grained transcriptomic approaches combined with thorough co-labelling techniques provide valuable insights into bivalve mollusk development

The developmental transcriptome of *M. galloprovincialis* presented in this study is currently unmatched in terms of sampling depth and detail with respect to the embryonic and larval stages represented in the dataset (Fig. 1). The sampling schedule in 4-hour intervals prevented biased sampling and overrepresentation of stereotypical stages (such as gastrula, trochophore or veliger), leading to the generation of a highly consistent dataset between biological replicates whose gene expression dynamics are dictated by developmental timing (Fig. 1B). The results of our differentially expression analysis, indicating a decrease in the number of DEGs as development proceeds, is seemingly in contrast with the findings of previous studies (Foulon et al., 2019; Moreira et al., 2018; Niu et al., 2016; Núñez-Acuña et al., 2022; Song et al., 2016a). However, it is important to note that differences in sampling strategies (*i.e.*, based on morphological criteria *versus* fixed time intervals) and settings of contrasts and thresholds in pairwise comparisons can yield different results. Our qPCR experiments corroborated the results obtained by our computational analyses, hence demonstrating the usefulness of our dataset for illustrating gene expression dynamics during early *M. galloprovincialis* development.

The representativeness of our transcriptome in the context of early bivalve development is further supported by the results of the qualitative analysis of the functional domains of DEGs. Transcription factors known to act in the regulation of metazoan development (*e.g.*, homeobox and fork head genes) and cohorts of genes involved in early shell formation were highly represented among the differentially expressed genes (Guo et al., 2023; Lartillot et al., 2002; Morino et al., 2013; Paps et al., 2015; Ramesh et al., 2019; Setiamarga et al., 2013; Zhao et al., 2018; Zheng et al., 2019). In addition, the analysis revealed a strong enrichment of receptors, transporters, and channels for hormones and neurotransmitters, which is consistent with previous analyses in bivalve mollusks implicating these mediators of intercellular signaling in the regulation of larval shell biogenesis, neurogenesis, and larval morphogenesis (Guo et al., 2023; Liu et al., 2020; Miglioli, 2022; Miglioli et al., 2021b; Song et al., 2016a). The fact that transmembrane transport and receptor-mediated signal transduction were highly significant biological processes in the *M. galloprovincialis* trochophore and advanced trochophore lends further support to this notion (Guo et al., 2023; Ramesh et al., 2019; Zhao et al., 2018; Zheng et al., 2019).

To correlate the transcriptional and developmental programs controlling early bivalve development, we set up a high-resolution, pan-larval protocol for multicolor *in situ* hybridization coupled with immunostaining in *M. galloprovincialis*. In combination with the developmental transcriptome, this protocol allows the parallel mapping of the expression dynamics of multiple genes in the context of cell type-specific markers. Using this approach, we were able to demonstrate the developmental expression and co-localization, in *M. galloprovincialis* embryos and larvae, of *Wnt8a* and *Foxb2,* which are key mediators of animal development, using immunoreactivity for the neurotransmitter serotonin (5HT) as a marker of the apical organ (Lartillot et al., 2002; Marlow et al., 2014; Miglioli et al., 2021a; Tomer et al., 2010). The combination of transcriptomic data and elaborate fluorescent whole-mount staining protocols has so far not been used for studying early development of bivalve mollusks and thus constitutes a major step towards the establishment of *M. galloprovincialis* as a model system.

### 3.2. Gene expression analyses suggest that the trochophore is a biphasic larval stage

Hierarchical clustering of libraries was carried out to identify developmental phases based on DEG dynamics (Leclère et al., 2019; Suzuki and Shimodaira, 2006). The results identified the main developmental phases of bivalve development (*i.e.*, zygote, gastrula, trochophore, and D-veliger) plus an additional phase, the advanced trochophore (Fig. 2A). In addition, a principal component analysis and a heatmap of Euclidean distances revealed a transcriptomic shift during the *M. galloprovincialis* trochophore stages, which is consistent with the notion that the trochophore is composed of two distinct developmental phases (Fig. 1). The existence of an advanced (or late) larval stage intercalated between the trochophore and the D-veliger has previously been proposed, but our results are the first to demonstrate the existence of this developmental phase on a transcriptomic level (Kamenev et al., 2018; Miglioli, 2022; Vogeler et al., 2016). Given that the morphotypes of the trochophore and of the advanced trochophore are very similar, our results indicate that developmental staging of bivalve mollusks by visual observation needs to be carried out with great care to avoid erroneous stage assignments. The existence of an additional larval stage linking the trochophore and the competent larva in mollusks, and possibly other lophotrochozoans, is an important discovery for evolutionary studies into the origin of the trochophore larva that sounds a note of caution for generalized comparisons of trochophores from distantly related species.

### 3.3. Identification of transcriptional transitions during bivalve mollusk development, with potential implications for shell formation and larval morphogenesis

Soft clustering revealed transcriptomic signatures overlapping the main developmental phases identified by hierarchical clustering. These signatures marked three developmental transitions: from gastrula to trochophore (Transition 1), from trochophore to advanced trochophore (Transition 2), and from advanced trochophore to D-veliger (Transition 3). Taking advantage of our fine-grained developmental sampling, the combined DEG and soft clustering analyses identified highly representative genes for each of the three transitions: 98 genes for Transition 1, 11 genes for Transition 2, and 8 genes for Transitions 3. The high number of Transition 1 genes marks the initiation of gene regulatory networks responsible for the ontogeny of the trochophore larva that includes developmental processes, such as neurogenesis, myogenesis, hematopoiesis, gut formation, and shell secretion (Arenas-Mena, 2013; Dyachuk and Odintsova, 2009; Dyrynda et al., 1997; Kniazkina and Dyachuk, 2022; Miglioli et al., 2019, 2021a; Song et al., 2016b; Yurchenko et al., 2018). In contrast, Transition 2 and Transition 3 are characterized by significantly lower numbers of genes, indicating that the transitions, respectively, of the trochophore to the advanced trochophore and of the advanced trochophore to the D-veliger transitions might require less dramatic transcriptomic changes.

To evaluate the potential evolutionary conservation of these three transitions within mollusks, we carried out a cross-species comparison with the Pacific oyster *C. gigas* (Liu et al., 2021). This comparison revealed that, while homologous sets of genes clearly marked Transition 1 (gastrula to trochophore) in the *C. gigas* developmental transcriptome, the conservation was much less apparent for Transition 2 (trochophore to advanced trochophore) and Transition 3 (advanced trochophore to D-veliger). It might be that inconsistent sampling designs of the two datasets blurred the transcriptional signatures of the transitions in *C. gigas*. Alternatively, the relative lack of conserved genes for Transition 2 and Transition 3 might be symptomatic of a global decrease of homologous transcriptomic traits during trochophore development of bivalve mollusks. In fact, phylostratigraphic analyses have revealed that the late trochophore stages of bivalves are characterized by the youngest transcriptome, which is largely derived from a class-specific phylostratum (Wu et al., 2019; Xu et al., 2016). Accordingly, specific adaptations of the larval body plan of bivalves, such as the shell field, originate from developmental programs initiated during the trochophore stage (Kemkemer and Long, 2014; Salamanca-Díaz et al., 2022; Xu et al., 2016).

To infer, which developmental processes might be controlled by conserved transition genes, we studied the expression of a subset of these genes in *M. galloprovincialis* embryos and larvae by *in situ* hybridization. For Transition 1 (16 to 20 hpf), we characterized the expression of 8 genes conserved between *M. galloprovincialis* and *C. gigas* (*Erg*, *Tbx2b*, *Dmbt1*, *Pxdn*, *SLC4A4*, *TYRO2*, *NA03*, and *HPP*) and found evidence for their involvement in developmental processes defining the transition from embryo to larva: ectomesoderm formation, dorsal specification, and shell field ontogeny. *Erg* and *Tbx2b* have respectively been described as pivotal regulators of ectomesoderm formation and dorsal specification in other lophotrochozoans, and the tissue localization in *M. galloprovincialis* early larvae is suggestive for a similar developmental role (Arenas-Mena, 2013; Osborne et al., 2018). With respect to shell field ontogeny, *Pxdn* and *TYRO2* are well known regulators of molluscan shell biogenesis, *SLC4A4* is an important player in shell calcification, and *Dmbt1* might be involved in the terminal differentiation of shell field columnar cells (Miglioli et al., 2019; Ramesh et al., 2019; Takito and Al-Awqati, 2004). The remaining two conserved genes of Transition 1 (*NA03* and *HPP*) could not be annotated, although similar sequences exist in other mollusks. However, we found expression of both genes in *M. galloprovincialis* to be associated with invaginating shell field tissue, which is suggestive of a potential role in larval shell biogenesis. For Transition 2 (28 to 32 hpf), we selected two genes conserved between *M. galloprovincialis* and *C. gigas*. Consistent with the notion that the period between 28 and 32 hpf marks the completion of a fully calcified larval shell in *M. galloprovincialis* (Miglioli et al., 2019), we found *SLC26A5*, a gene encoding a bicarbonate transporter that provides substrate for shell calcification (Ramesh et al., 2019), expressed in two symmetric groups of ectodermal cells at Transition 2. The second gene, *Grin3a*, encodes the NMDA-type receptor subunit NR3, and, at Transition 2, it was most conspicuously expressed in a group of cells close to the apical tuft. NMDA subunits NR1 and NR2 have recently been characterized in nerves and ganglions of *C. gigas,* but the functions of NMDA-type receptors during bivalve trochophore development have yet to be assessed (Vogeler et al., 2021).

Transition 3 (40 to 44 hpf) marks the internalization of the velum into the D-shaped shell, which requires a functional muscular system (Dyachuk and Odintsova, 2009). The first of the two conserved transition genes we selected for Transition 3, *EF-hand*, has been linked to myosin regulation in scallop muscle fibers, and the expression in *M. galloprovincialis* D-veligers shows similarities to the ventral distribution of paramyosin immunoreactivity in D-veliger larvae of the mussel *M. trossulus* (Dyachuk and Odintsova, 2009; Fromherz and Szent-Györgyi, 1995). The second gene, *COLVI*, is expressed in mantle cells of *M. galloprovincialis*, on both sides of the D-border of the larval shell, which is consistent with a function in nacre formation, as has previously been suggested for the *COLVI* homolog of the pearl oyster *Pinctada martensii* (Zheng et al., 2017).

Taken together, the *in situ* hybridization-based expression analysis of conserved transition genes highlighted developmental processes that might be regulated by homologous sets of genes in *M. galloprovincialis* and *C. gigas*. We thus identified genes with possible functions in ectomesoderm formation, dorsal specification, muscle development as well as shell ontogeny, secretion, and calcification, whose expression is likely characteristic of bivalve mollusk development. We hypothesize that the transcriptomic staging approach we used for the characterization of *M. galloprovincialis* development can be applied to other species (bivalves, mollusks, lophotrochozoans) and used in cross-species comparisons for (1) the definition of developmental stages at a transcriptomic level, (2) the identification of major developmental transitions in transcriptome datasets, and (3) the comparison of homologous gene sets characteristic of a given developmental phase or transition.

### 3.4. The Mediterranean mussel, *Mytilus galloprovincialis*, a novel model for developmental studies of mollusks

The results reported in this study represent the first thorough transcriptomic analysis of the development of the Mediterranean mussel *M. galloprovincialis*. It further presents a collection of protocols essential for the validation and exploitation of transcriptomic data, including experimental tools, such as qPCR, immunohistochemistry, and *in situ* hybridization. The combination of fluorescent immunohistochemistry and *in situ* hybridization is particularly powerful for assessing the co-localization of crucial developmental factors in mussel embryos and larvae. Potential applications for this transcriptome, and associated experimental protocols, include developmental studies, cross-species comparisons, and ecotoxicological surveys. Taking advantage of the developmental transcriptome and the novel protocols, we are the first to document the expression of a suite of genes in *M. galloprovincialis* that characterize bivalve mollusk development, revealing the full potential of this animal system for becoming a reference for developmental studies using mollusk models.

## 4. Materials and Methods

### 4.1. Spawning, fertilization, and rearing of embryos and larvae

Sexually mature *M. galloprovincialis* adults were harvested from the natural population in the bay of Villefranche-sur-Mer, France (43.682°N, 7.319°E) during the spawning seasons of 2021, 2022, and 2023 (January through April). Animals were acclimated to and maintained in laboratory conditions, as previously described (Miglioli et al., 2021b). Thirty adult mussels were exposed to a heat shock to induce gamete release. Briefly, epibionts, dirt, and algae were scrubbed off the shells under running filtered natural seawater. Byssus and threads were carefully cut off. Clean mussels were placed on ice for 10 minutes and then immersed, in individual 200 mL containers and on a rocking shaker, in filtered natural seawater kept at 28°C until gamete release. Spawning mussels were immediately moved to filtered natural seawater at 16°C. Prior to fertilization, gametes were collected, washed, and maintained in Millipore-filtered natural seawater (using a 0.2 µm Millipore filter). Fertilization was carried out as previously described with a 1:10 ratio of eggs to sperm (Miglioli et al., 2021a, 2021b). After 30 minutes, the fertilization success rate (*i.e.*, the ratio of the number of fertilized eggs over the total number eggs x 100) was established by microscopic observation. Parental pairs with a high fertilization success rate (> 90%) were selected for the following experimental manipulations. Two experimental replicates were set up, each consisting of a homogeneous pool of fertilized eggs from two independent parental pairs. Embryos from each replicate were distributed into 200 mL flasks for suspended cultures and brought to a density of 100 embryos per mL with Millipore-filtered natural seawater. A total of 14 flasks per replicate were prepared, corresponding to the pre-defined sampling strategy: in addition to sampling before fertilization, sample collections were carried out every 4 hours from 4 hpf to 52 hpf, with an additional sampling timepoint at 72 hpf. Embryo cultures were maintained at 16°C throughout the experiment. Samples for immunohistochemistry and *in situ* hybridization were produced following the same culture protocol. After filtration, larvae were fixed in 4% paraformaldehyde in 1x PBS overnight at 4°C, washed 3 times for 15 minutes in 1x PBS with 0.01% Tween 20, and then stored in 100% methanol at -20°C (Miglioli et al., 2021b).

### 4.2. Sampling, total RNA extraction, and high-throughput sequencing

RNA extraction and purification were carried out with the RNAqueous-Micro Kit (Invitrogen) and the RNeasy MinElute Cleanup Kit (Qiagen) following published protocols (Leclère et al., 2019). Unfertilized eggs were directly collected from spawning females, and embryos and larvae were sampled with a 60 µm mesh filter and concentrated in RNAse free 1.5 mL tubes. The tubes were then placed on ice and gently centrifuged multiple times to remove the seawater from the samples. The pellet was immediately resuspended in lysis buffer, homogenized by energetic pipetting, and flash-frozen in liquid nitrogen. For each sample, 10 µL of the 1.5 mL filtrate were placed in a separate tube and fixed overnight in 4% paraformaldehyde in 1x PBS at 4°C (Miglioli et al., 2021a). The fixed larvae were stained with Hoechst (Invitrogen) and imaged with a SP8 confocal microscope (Leica Microsystems) to validate proper developmental progression (Miglioli et al., 2021a). Lysed samples were stored at -80 °C until RNA extraction and purification (Leclère et al., 2019). RNA concentration and quality was assessed with a Nanodrop (Thermo Scientific) and an Agilent 2100 Bioanalyzer. Samples were shipped on dry ice to BGI (Hong Kong) for high-throughput sequencing. At least thirty million clean paired-end reads of 200 bp each were produced using the BGISEQ-500 Platform. Read cleaning and trimming was performed by BGI (Hong Kong). After sequencing, the raw reads were filtered. Data filtering included removal of adapter sequences, contaminations, and low-quality reads from the raw data. The entire read was deleted, if more than 28% matched the adapter sequence, if more than 40% had a quality value lower than 20 or if there were more than 3% of unidentifiable nucleotides in the read. This step was completed by the SOAP nuke software developed by BGI. Libraries were not subjected to further trimming (Dobin et al., 2013).

### 4.3. Genome-guided transcriptome assembly and differential expression analysis

Sequencing quality of the libraries was assessed with FastQC (v. 0.11.7) (Andrews, 2010). The libraries were mapped on the indexed *M. galloprovincialis* reference genome with STAR (v. 2.7.10a) using default parameters (Dobin et al., 2013; Gerdol et al., 2020). Read counts were obtained using StringTie (v2.2.1) in counting mode with the genome annotation file (Pertea et al., 2015, 2016). The resulting gene count matrix was exported using prepDE.py and processed in Rstudio (v. 4.2.2) with the DESeq2 (v. 1.24.0) package using default parameters and a design file indicating replicate number and developmental time for each library (Love et al., 2014). The dataset was then pre-filtered by keeping all genes with at least 10 reads in at least two replicated libraries to reduce the potential batch effect derived from the variability in gene presence-absence characterizing the species (Gerdol et al., 2020). The resulting matrix was normalized using the regularized logarithms of counts (DESeq2::rlog) and used to perform principal component analyses and heatmaps (Kolde, 2019). Stage-by-stage differential expression analysis was performed with DESeq2 using the pairwise comparison function, with the reference stage for comparisons relevelled for each contrast (Love et al., 2014). Pairwise comparisons were performed on samples of consecutive stages (0 hpf *versus* 4 hpf, 4 hpf *versus* 8 hpf, etc.). DEGs were filtered by adjusted p-value (padj.cutoff < 0.05) and log2 fold change (lfc.cutoff > 0.58).

### 4.4. Developmental staging and gene clustering

To discriminate stages of *M. galloprovincialis* development from a transcriptomic perspective, bootstrapped hierarchical clustering with pvclust (v. 2.2-0) default parameters was performed on regularized logarithms (rlog) of DEG counts as previously described (Leclère et al., 2019; Love et al., 2014; Suzuki and Shimodaira, 2006). DEGs (rlog normalized counts) were then clustered with DegReport (v. 1.36.0) using default parameters to identify groups of DEGs with similar expression dynamics during *M. galloprovincialis* development (Pantano, 2023). Soft clustering with the Mfuzz (v. 2.56.0) package was performed on the transcripts per million normalized matrix to identify representative gene clusters of different developmental phases and potential transcriptomic waves (Futschik and Carlisle, 2005). The number of soft clusters was determined using the elbow method to the minimum centroid distance (Futschik and Carlisle, 2005). Genes were assigned to the cluster with the highest membership probability.

### 4.5. Functional annotation, gene ontologies, and enrichment analyses

The predicted protein sequences of the DEGs were annotated using BLASTp (v. 2.11.0) against the UniProt/SwissProt database, conserved domains were identified with hmmerscan from HMMER 3.3 against the Pfam database, and transcription factor domains were screened with AnimalTFDB (v2.0) (Finn et al., 2015, 2017; Jones et al., 2014; Mahram and Herbordt, 2015; Sayers et al., 2021; Zhang et al., 2015; Mistry et al., 2021). Ontology enrichment analysis was performed on the list of DEGs per stage and soft cluster. The protein sequence corresponding to genes of interest were extracted from the *M. galloprovincialis* proteome and annotated with Gene Ontology (GO) identifiers by Interproscan (v. 95.0) analysis (Finn et al., 2017; Gerdol et al., 2020; Jones et al., 2014). Enrichment analysis of Biological Processes (BP) was conducted with topGO R (v3.17). Significance of the enrichment was computed with Fisher’s exact test (Alexa and Rahnenführer, 2023), and significant lists of GOs were clustered by k-means using semantic similarity of SimplifyEnrichment (v. 1.2.0) to identify potential sets of genes responsible for stage-enriched BPs (Gu and Hübschmann, 2022).

### 4.6. Transcriptome validation by real time quantitative polymerase chain reaction

Complementary first strand DNAs (cDNAs) were synthesized from total RNA extracts of both biological replicates at all sample timepoints using the SuperScript VILO Kit (Invitrogen). cDNA synthesis was performed using random primers following the manufacturer’s instructions: 10 minutes at 25°C, 60 minutes at 42°C, and 5 minutes at 85°C. To validate the differential gene expression analysis and the cluster analysis, four target genes were selected: *Wnt8a* (MGAL_10B085403), *Foxb2* (MGAL_10B093191), *SLC46A3* (MGAL_10B020966), and *SYTL4* (MGAL_10B044489). *EF-*α*1* was used for data normalization as one of the best-performing reference gene products (Balbi et al., 2016). Primer pairs were designed within the open reading frame of each gene using Primer3 (Koressaar and Rem, 2007). Sequences, efficiency, and amplicon sizes are available in the supplementary material (Supplementary Table 9). qPCR reactions were carried out for each set of primers, in experimental triplicates. The amplification was followed in real time using LightCycler 480 SYBR Green 1 master mix (Roche) in 45 cycles (initial 5 minutes at 95°C, followed by 45 cycles of 20 seconds at 95°C, 20 seconds at 60°C, and 30 seconds at 72°C) in a LightCycler 480 (Roche). Quantification was carried out in parallel for each of the target genes and the reference gene. The quantification of the expression at a given stage was determined in relation to the expression of the reference gene using the comparative CT method (Schmittgen and Livak, 2008). To compare qPCR and transcriptome results, expression of the target genes from both datasets were normalized with respect to their maximum value. Statistical differences in qPCR results were assessed using a Student’s t-test (de Winter, 2013).

### 4.7. *In situ* hybridization using hybridization chain reaction and co-labeling by immunohistochemistry

Hybridization probe sets were designed using a user-friendly python interface (https://github.com/rwnull/insitu_probe_generator) and synthesized by OligoPools (Twist Bioscience). Amplifiers in red (B2: 546 nm) and far red (B1: 647 nm) were purchased from Molecular Instruments (https://www.molecularinstruments.com/). All probe sets with relative amplifier pairings are reported in Supplementary Table 10. The larval samples previously collected and stored in methanol were re-hydrated with four washes of 5 minutes followed by a 10-minute incubation in 1x PBS with 0.01% Triton X-100 at room temperature. Permeabilization was carried out by a 30-minute incubation in the detergent solution (1% SDS, 0.005% Tween 20, 50 mM HCl pH 7.5, 1 mM EDTA, pH 8, 0.15 M NaCl) at room temperature. After two washes of 1 minute and two washes of 5 minutes in 1x PBS with 0.01% Triton X-100 at room temperature, the specimens were placed in pre-hybridization for 30 minutes at 37°C with gentle shaking in hybridization buffer (HB) (30% formamide, 5x SSC, 9 mM citric acid, pH 6, 0.01% Tween 20, 2 mg heparin, 1x Denhardt’s solution, 10% dextran sulphate). After addition of the probes (at a final concentration of 4 mM), hybridization was carried out overnight. Subsequent washes were performed at 37°C: starting with an initial addition of 1.5 volumes of washing solution (30% formamide, 5x SSC, 9 mM citric acid, pH 6, 0.01% Tween 20, 2 mg Heparin) with respect to the volume of the hybridization solution, four washes of 15 minutes were performed with washing solution. Specimens were then washed, at room temperature, twice for 5 minutes in 5x SSC with 0.01% Tween 20, once for 5 minutes in 1x SSC with 0.01% Tween 20, and 30 minutes in amplification buffer (5x SSC, 0.01% Tween 20, 10% dextran sulphate), before amplifiers were added (at a final concentration of 60 mM) to start the amplification chain reaction. Amplifiers were prepared as follows: the required volume of hairpins 1 and 2 from each amplifier were placed in a 0.6 ml tube and incubated for 90 seconds at 95°C. After the heat shock, amplifiers were placed in the dark at room temperature for 30 minutes. The amplifiers were then resuspended in amplification buffer and distributed to the samples. Amplification was carried out at room temperature in the dark overnight. Excess hairpins were removed in two washes of 5 minutes and two washes of 30 minutes in 1x SSC with 0.01% Tween 20, followed by two washes of 5 minutes in 1x PBS with 0.01% Tween 20. Co-labeling by immunocytochemistry with serotonin (5-HT) anti-rabbit antibody (1:10,000; #20080 Immunostar, Hudson, WI, USA) and goat anti-rabbit Alexa Fluor 488 conjugated secondary antibody (1:500; #4412S Cell Signaling Technology, Leiden, The Netherlands) was performed immediately after HCR amplification using previously described conditions (Miglioli et al., 2021a, 2021b). Specimens were then prepared for imaging by incubation in 1 µg/L Hoechst (Invitrogen) in 1x PBS for 30 minutes at room temperature. Following three washes of 5 minutes in 1x PBS with 0.01% Tween 20, specimens were mounted with the antifading agent CitiFluor AF1 (Agar Scientific) and imaged on a SP8 confocal microscope (Leica Microsystems), as previously described (Miglioli et al., 2021b). Images were analyzed with ImageJ (Schneider et al., 2012) and fluorescent signals were corrected as follows: autofluorescence from tissue and shell was subtracted with “math”, the signal was homogenized with the “despeckle” or “smooth” functions, and z-stacks were merged by either “maximum intensity” or “sum slices”.

### 4.8. Developmental transition gene selection and comparisons with the developmental transcriptome of the Pacific oyster *Crassostrea gigas*

Representative genes of the developmental transitions were identified in *M. galloprovincialis* following this strategy. Developmental transitions were defined based on the hierarchical clustering of the stages. Three transitions were identified: from 16 to 20 hpf, 28 to 32 hpf, and 40 to 44 hpf. Genes were subsequently clustered according to their expression profile by soft clustering (see section 4.4), allowing the identification of three gene clusters corresponding to the three transitions. From these clusters, only genes showing a differential expression at the time of transition were selected. For Transition 1, genes were selected, if they were differentially expressed (log fold change ≥ 2, approximated by excess) between 16 and 20 hpf. For Transition 2, genes were selected, if they were differentially expressed (log fold change ≥ 2, approximated by excess) between 28 and 32 hpf. For Transition 3, genes were selected, if they were differentially expressed (log fold change ≥ 2, approximated by excess) between 40 and 44 hpf. To identify potentially conserved genes, a BLASTp search (Mahram and Herbordt, 2015) of the *M. galloprovincialis* transition genes was performed against the proteome of the Pacific oyster *C. gigas* (Liu et al., 2021). Their expression profiles in *C. gigas* was then visualized with a heatmap (Kolde, 2019) to investigate the presence of similar developmental transitions in *M. galloprovincialis* and *C. gigas*. *In situ* hybridization experiments were subsequently carried out on a selection of the conserved transition genes in *M. galloprovincialis* (see section 4.7). All selected *M. galloprovincialis* genes had a homolog in *C. gigas*, whose z-score value turned positive (> 0) at the time of the transition.

## 5. Acknowledgments

The authors are indebted to Guy Lhomond and Lucas Leclère for technical assistance and fruitful discussions. The authors would like to thank Sébastien Schaub of the Plateforme d’Imagerie par Microscopie (PIM) as well as Laurent Gilletta and Axel Duchene of the Service Aquariologie (SA) of the Institut de la Mer de Villefranche (IMEV), France, that is supported by EMBRC-France and funded by the Agence Nationale de la Recherche (ANR) (ANR-10-INBS-02). We thank the members of the Ascidian BioCell and EvoInSiDe teams at the Laboratoire de Biologie du Développement de Villefranche-sur-Mer, France, as well as the members of the team of Environmental Physiology at the Università Degli Studi Di Genova, Italy, for their support of this work.

## 8. Data Availability

All libraries analyzed in this article have been deposited in the NCBI SRA database (Accession: PRJNA996031; ID: 996031).

## 11. Supplementary Figure Legends

**Supplementary Figure 1:**
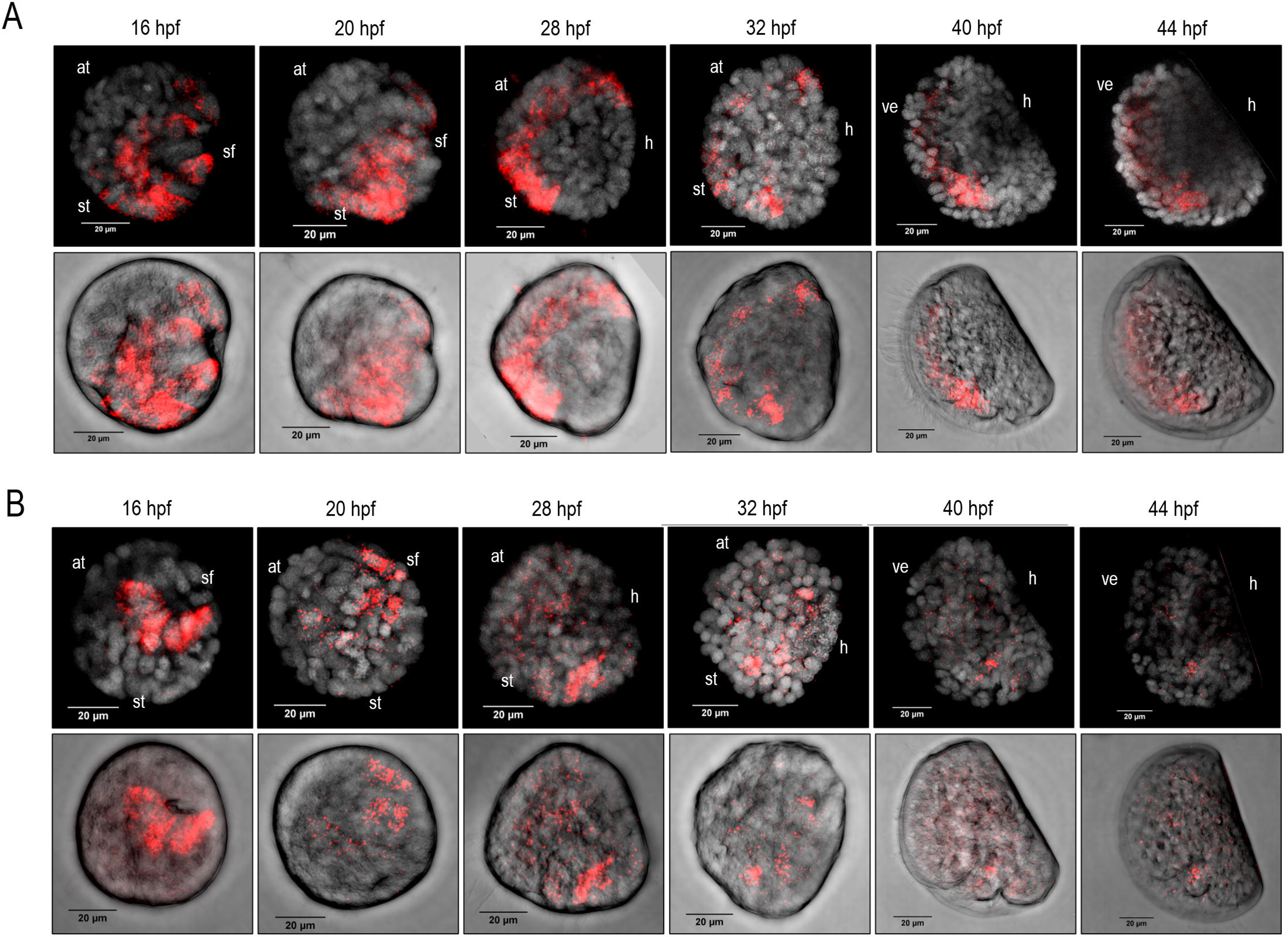
Fluorescent *in situ* hybridization protocol validation with *Foxb2* and *Wnt8a* probes at different stages of *Mytilus galloprovincialis* development. A) Expression pattern of *Foxb2*. Upper panel shows maximum z-projections of the *in situ* hybridization signal (red) with Hoechst nuclear staining (grey). Lower panel shows the *in situ* hybridization signal (red) in brightfield. B) Expression pattern of *Wnt8a*. Upper panel shows maximum z-projections of the *in situ* hybridization signal (red) with Hoechst nuclear staining (grey). Lower panel shows the *in situ* hybridization signal (red) in brightfield. Abbreviations: at: apical tuft; h: hinge; sf: shell field; st: stomodeum/presumptive mouth; ve: velum. Scale bar: 20μm.

**Supplementary Figure 2:**
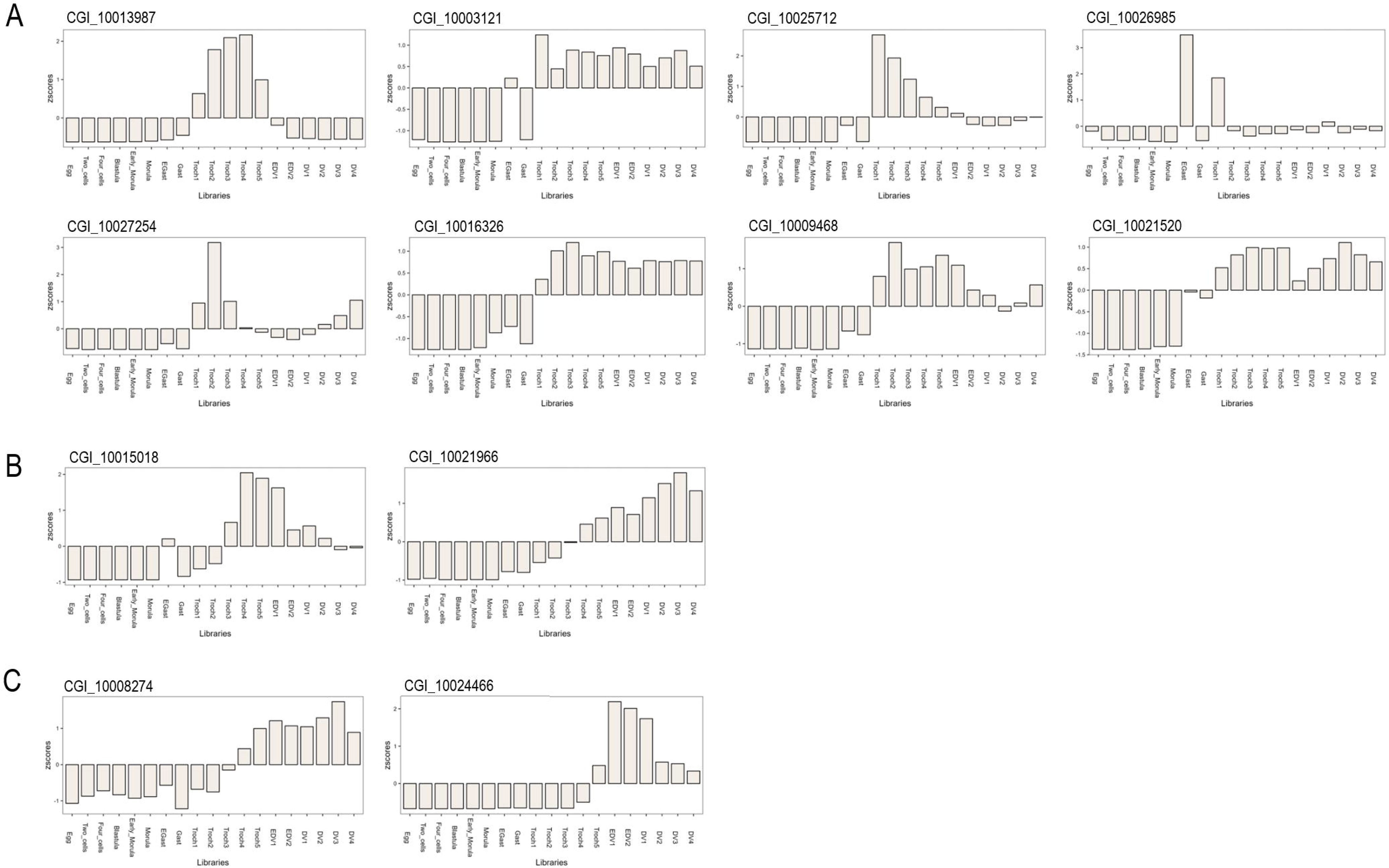
Developmental expression patterns of transition genes in a developmental transcriptome of the Pacific oyster *Crassostrea gigas*. A) Transition1. B) Transition 2. C) Transition 3.

## 12. Supplementary Files and Tables

***Supplementary File 1:*** Supplementary Table 1. Summary of mapping results for each library. Supplementary Table 2. Matrix of raw gene counts. Supplementary Table 3. Dynamics of differentially expressed genes with Uniprot protein annotation, Pfam domain annotation, DEG cluster annotation, developmental transition membership, and soft clustering membership. Supplementary Table 4. DegReport analysis. The mean expression value of each cluster is reported for each timepoint, and the expression trend is presented as a sparkline histogram. The green bar indicates the highest and the red one the lowest value. Supplementary Table 5. Binary matrix of the DegReport output. Supplementary Table 6. List of differentially expressed transcription factors identified with AnimalTFDB. Supplementary Table 7. Binary matrix of the Mfuzz soft clustering analysis. Supplementary Table 8. Best blastp hits of *Mytilus galloprovincialis* transition genes in the *Crassostrea gigas* genome. Supplementary Table 9. Primer pairs used for qPCR analyses. Supplementary Table 10. Probe sets for *in situ* hybridization by hybridization chain reaction used in the study.

***Supplementary File 2:*** <u>DESeq2 output of differentially expressed genes by pairwise</u> <u>comparison</u>. Every tab corresponds to a pairwise comparison.

